## Supporting Information for "Dynamic Architecture of Mycobacterial Outer Membranes Revealed by All-Atom Simulations"

**Table S1. System Compositions and Basic Statistics†.**

| System Name (_Temp) | Leaflet | MA_eU : MA_sZ : MA_W : PDIM : TDM : TMM : DAT : PAT : SGL | System Size (Å) | Membrane Thickness (Å) | APL (Å$^2$) |
|---|---|---|---|---|---|
| MA_eU_313 | both | **60 : 00 : 00 : 00 : 00 : 00 : 00 : 00 : 00** | 62.4 | 59.4 | 64.9 |
| MA_eU_323 | both | **60 : 00 : 00 : 00 : 00 : 00 : 00 : 00 : 00** | 63.2 | 58.8 | 66.5 |
| MA_eU_333 | both | **60 : 00 : 00 : 00 : 00 : 00 : 00 : 00 : 00** | 63.6 | 9.0 | 67.5 |
| MA_eU_338 | both | **60 : 00 : 00 : 00 : 00 : 00 : 00 : 00 : 00** | 65.0 | 59.4 | 70.3 |
| MA_eU_343 | both | **60 : 00 : 00 : 00 : 00 : 00 : 00 : 00 : 00** | 66.0 | 59.2 | 72.6 |
| MA_eU_353 | both | **60 : 00 : 00 : 00 : 00 : 00 : 00 : 00 : 00** | 67.2 | 57.8 | 75.3 |
| MA_sZ_313 | both | **00 : 60 : 00 : 00 : 00 : 00 : 00 : 00 : 00** | 60.2 | 65.0 | 60.3 |
| MA_W_313 | both | **00 : 00 : 60 : 00 : 00 : 00 : 00 : 00 : 00** | 58.6 | 70.8 | 57.3 |
| MA_eU_W_313 | both | **30 : 00 : 30 : 00 : 00 : 00 : 00 : 00 : 00** | 61.9 | 61.7 | 63.9 |
| MA_sZ_333 | both | **00 : 60 : 00 : 00 : 00 : 00 : 00 : 00 : 00** | 64.4 | 61.2 | 69.2 |
| MA_W_333 | both | **00 : 00 : 60 : 00 : 00 : 00 : 00 : 00 : 00** | 64.4 | 61.4 | 69.0 |
| MA_eU_W_333 | both | **30 : 00 : 30 : 00 : 00 : 00 : 00 : 00 : 00** | 64.7 | 60.8 | 69.8 |
| All_Lipids_313 | both | **00 : 00 : 00 : 20 : 20 : 20 : 20 : 20 : 20** | 109.3 | 56.3 | 99.6 |
| No_PDIM_313 | both | **00 : 00 : 00 : 00 : 24 : 24 : 24 : 24 : 24** | 117.6 | 49.1 | 115.2 |
| No_TDM_313 | both | **00 : 00 : 00 : 24 : 00 : 24 : 24 : 24 : 24** | 106.0 | 53.8 | 93.6 |
| No_TMM_313 | both | **00 : 00 : 00 : 24 : 24 : 00 : 24 : 24 : 24** | 110.0 | 57.3 | 100.8 |
| No_DAT_313 | both | **00 : 00 : 00 : 24 : 24 : 24 : 00 : 24 : 24** | 113.7 | 57.4 | 107.7 |
| No_PAT_313 | both | **00 : 00 : 00 : 24 : 24 : 24 : 24 : 00 : 24** | 105.4 | 57.3 | 92.6 |
| No_SGL_313 | both | **00 : 00 : 00 : 24 : 24 : 24 : 24 : 24 : 00** | 102.8 | 60.1 | 88.0 |
| All_Lipids_333 | both | **00 : 00 : 00 : 20 : 20 : 20 : 20 : 20 : 20** | 106.3 | 59.6 | 94.1 |
| No_PDIM_333 | both | **00 : 00 : 00 : 00 : 24 : 24 : 24 : 24 : 24** | 113.0 | 54.5 | 106.3 |
| No_TDM_333 | both | **00 : 00 : 00 : 24 : 00 : 24 : 24 : 24 : 24** | 101.8 | 60.9 | 86.4 |
| No_TMM_333 | both | **00 : 00 : 00 : 24 : 24 : 00 : 24 : 24 : 24** | 106.7 | 62.0 | 94.8 |
| No_DAT_333 | both | **00 : 00 : 00 : 24 : 24 : 24 : 00 : 24 : 24** | 109.4 | 61.9 | 99.8 |
| No_PAT_333 | both | **00 : 00 : 00 : 24 : 24 : 24 : 24 : 00 : 24** | 105.5 | 56.5 | 92.8 |
| No_SGL_333 | both | **00 : 00 : 00 : 24 : 24 : 24 : 24 : 24 : 00** | 101.0 | 64.2 | 84.0 |
| Asym_313 | inner | **94 : 47 : 47 : 00 : 00 : 00 : 00 : 00 : 00** | 110.1 | 61.0 | 59.0 |
| | outer | **00 : 00 : 00 : 20 : 20 : 20 : 20 : 20 : 20** | | | 92.4 |
| Asym_333 | inner | **94 : 47 : 47 : 00 : 00 : 00 : 00 : 00 : 00** | 105.3 | 63.8 | 64.5 |
| | outer | **00 : 00 : 00 : 20 : 20 : 20 : 20 : 20 : 20** | | | 101.1 |

†The system size, membrane thickness, and area per lipid (APL) are the averaged numbers of the last 500 ns production.

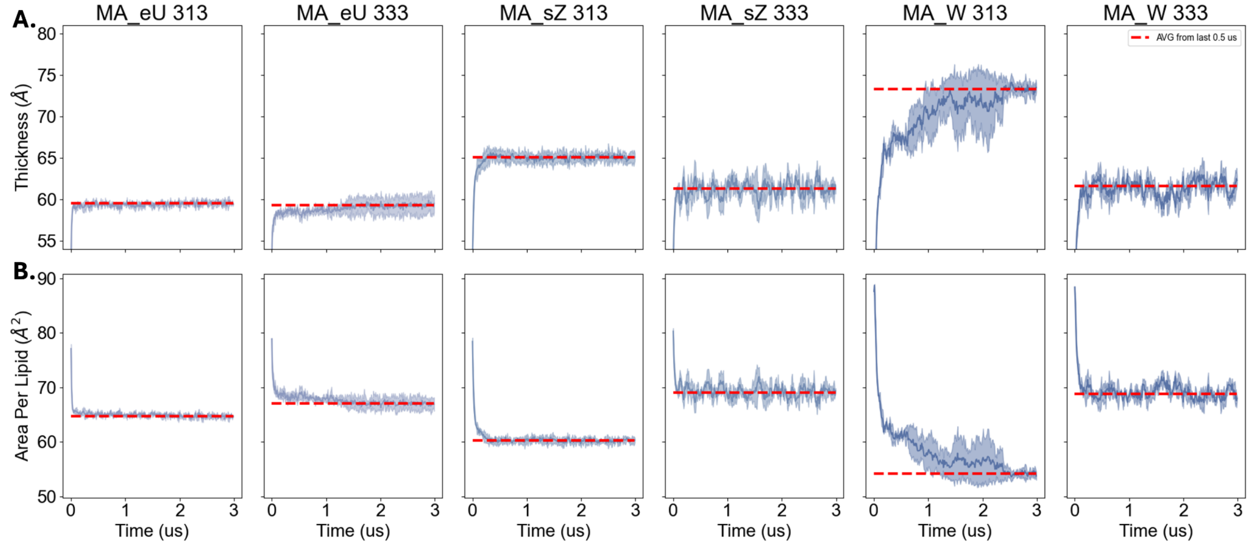


**Figure S1. Time Series of Inner Leaflet Symmetric Bilayers with Various Starting Conformations of α-mycolic acid. A.** Membrane thickness. **B.** Area per lipid.

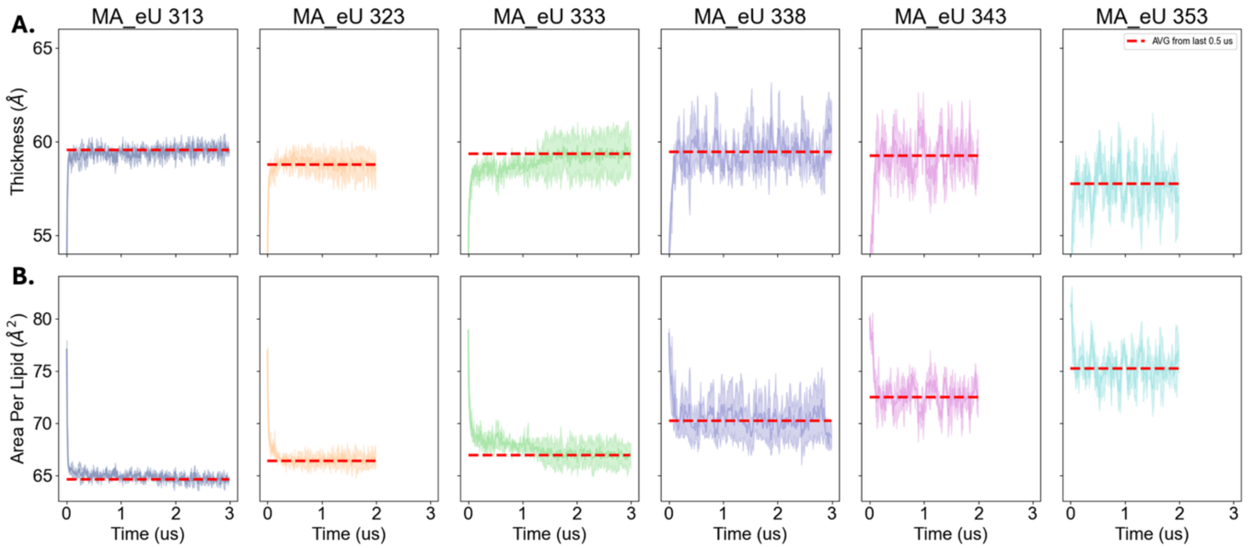


**Figure S2. Time Series of MA_eU Bilayers at Increasing Temperature**. **A.** Membrane thickness. **B.** Area per lipid.

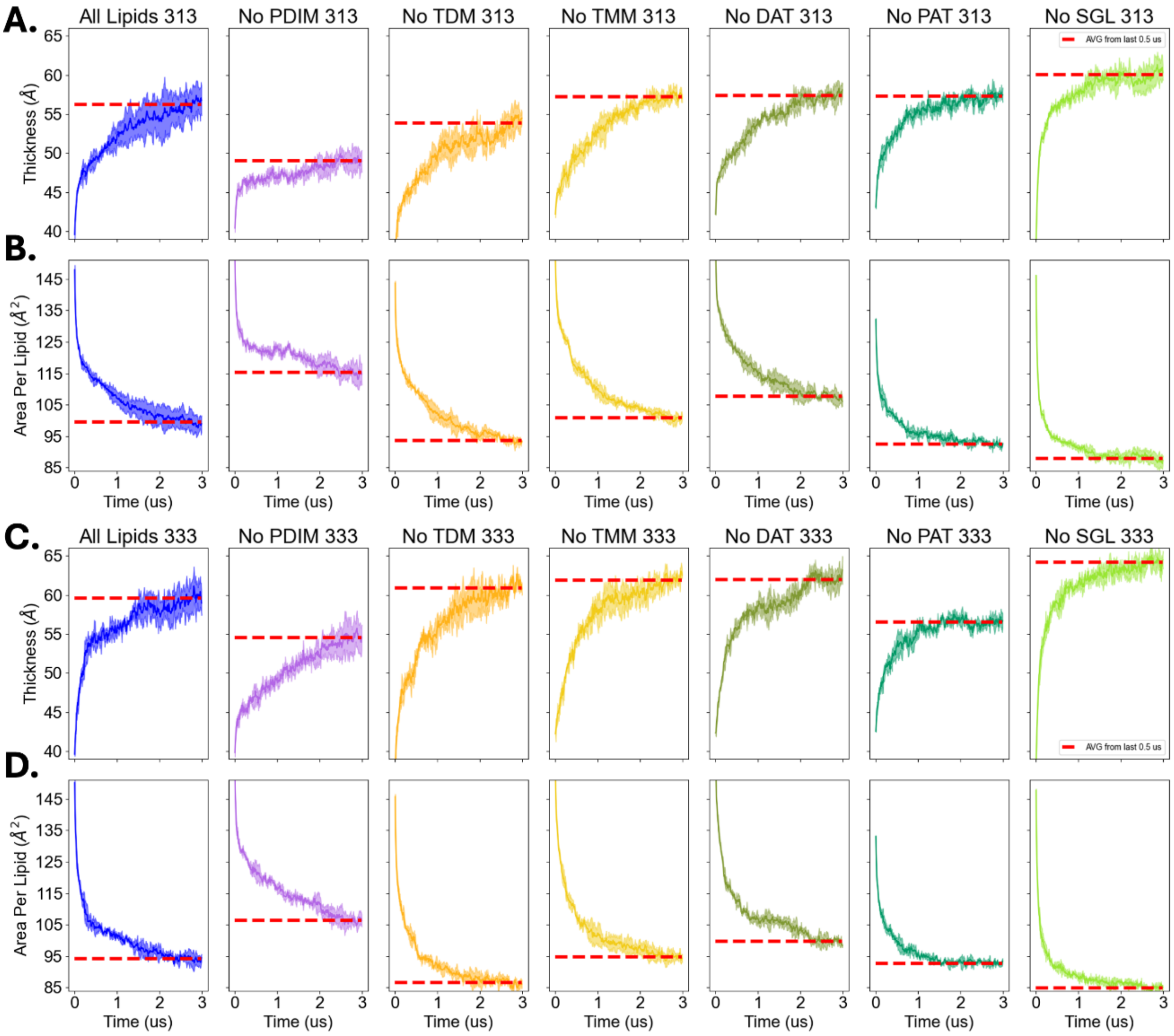


**Figure S3. Time Series of Outer Leaflet Symmetric Systems at 313K and 333K**. **A,C.** Membrane thickness. **B,D.** Area per lipid.

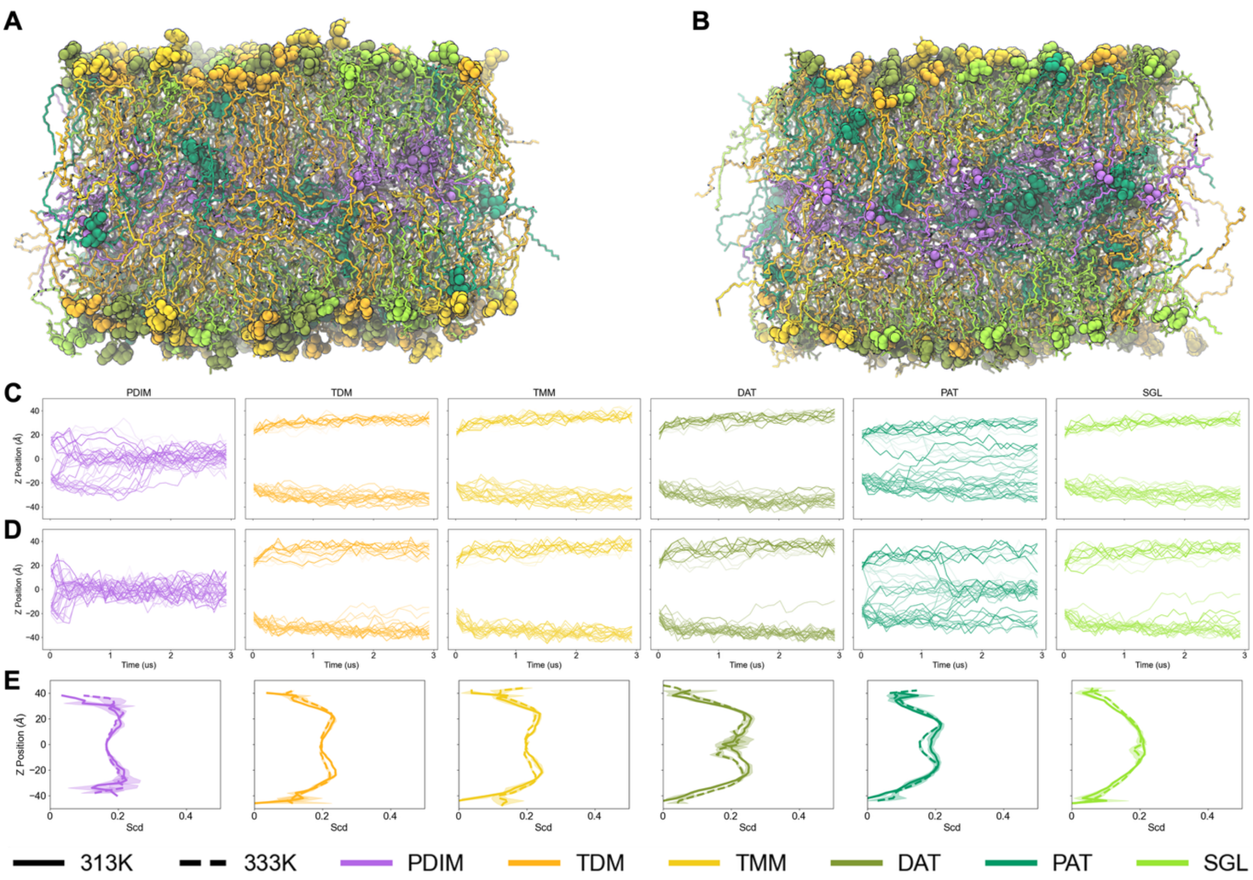


**Figure S4. Outer Leaflet Lipid Dynamics. A,B.** Snapshots of All_Lipids system after 3 μs at 313 K and 333 K, respectively. Lipid headgroups are shown as van der Waals spheres, and the rest of each lipid is shown with the QuickSurf drawing method. **C,D.** Headgroup z position time series of all lipid types at 313K and 333K, respectively. **E.** Average deuterium order parameters of carbons along the z-axis for all lipid types. Solid lines are 313 K and dashed lines are 333 K. Scd was averaged over the final 500 ns production.

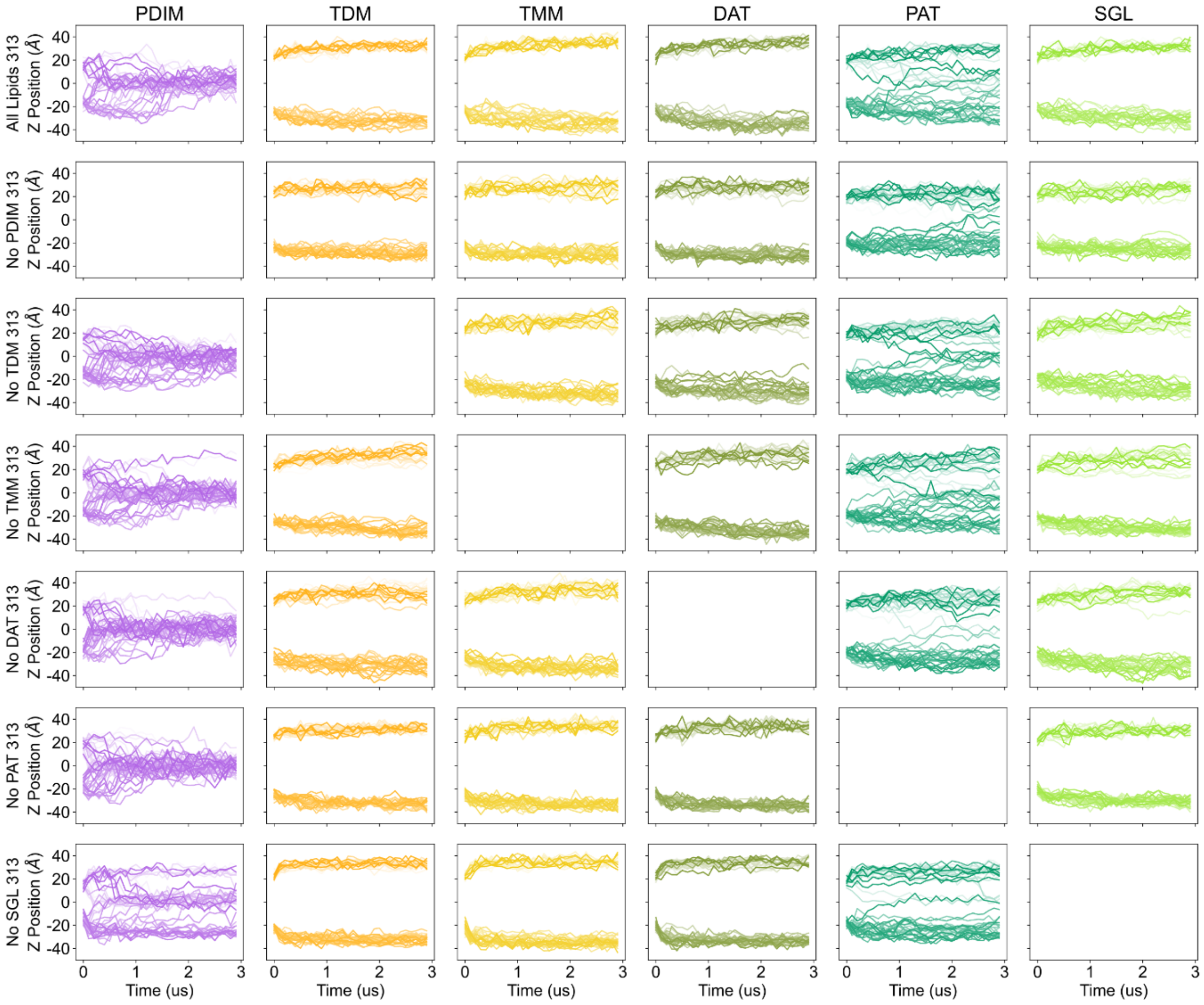


**Figure S5. Headgroup Z Position Time Series for All Lipid Types and All Outer Leaflet Symmetric Systems at 313K.**

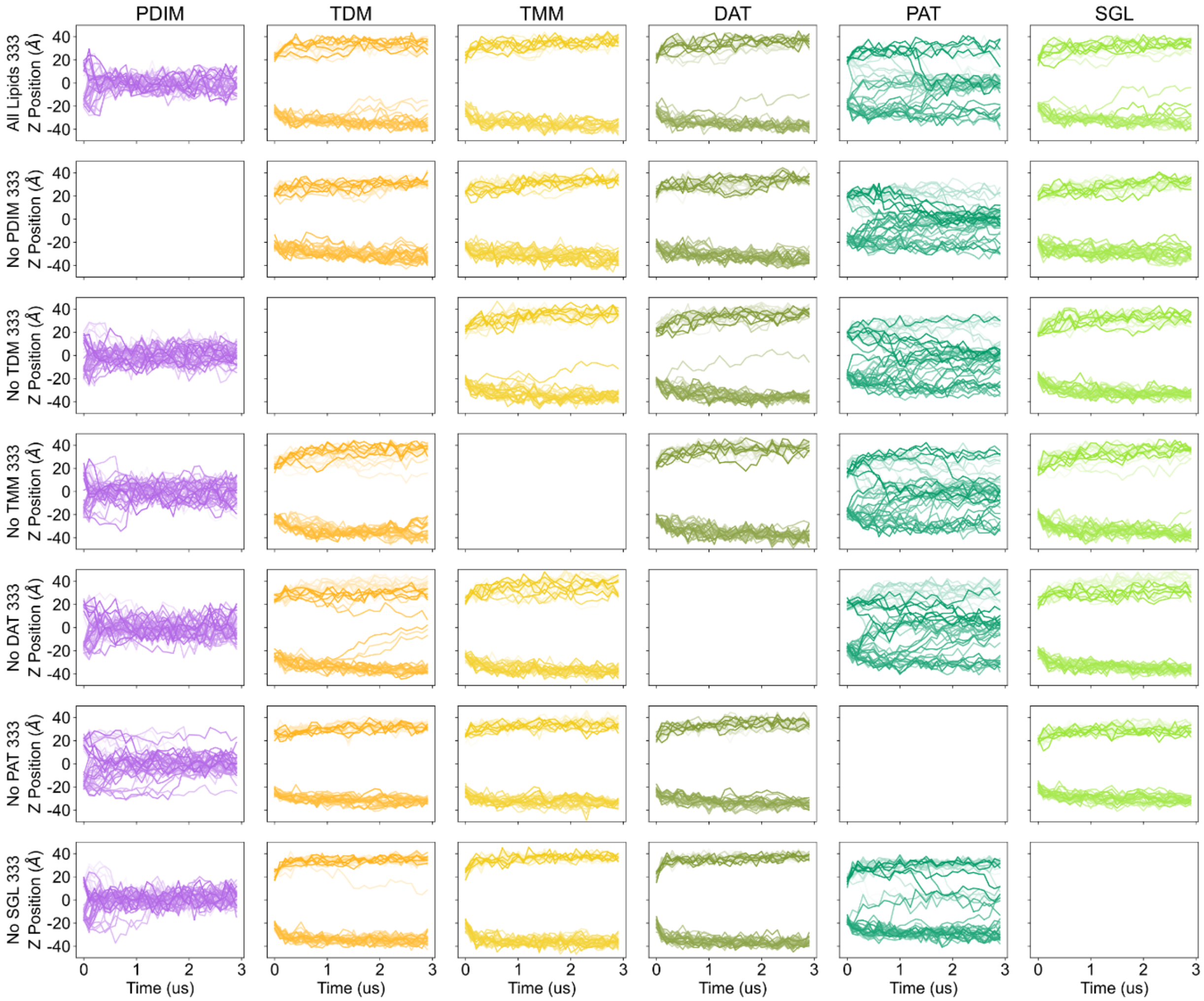


**Figure S6. Headgroup z Position Time Series for All Lipid Types and All Outer Leaflet Symmetric Systems at 333K.**

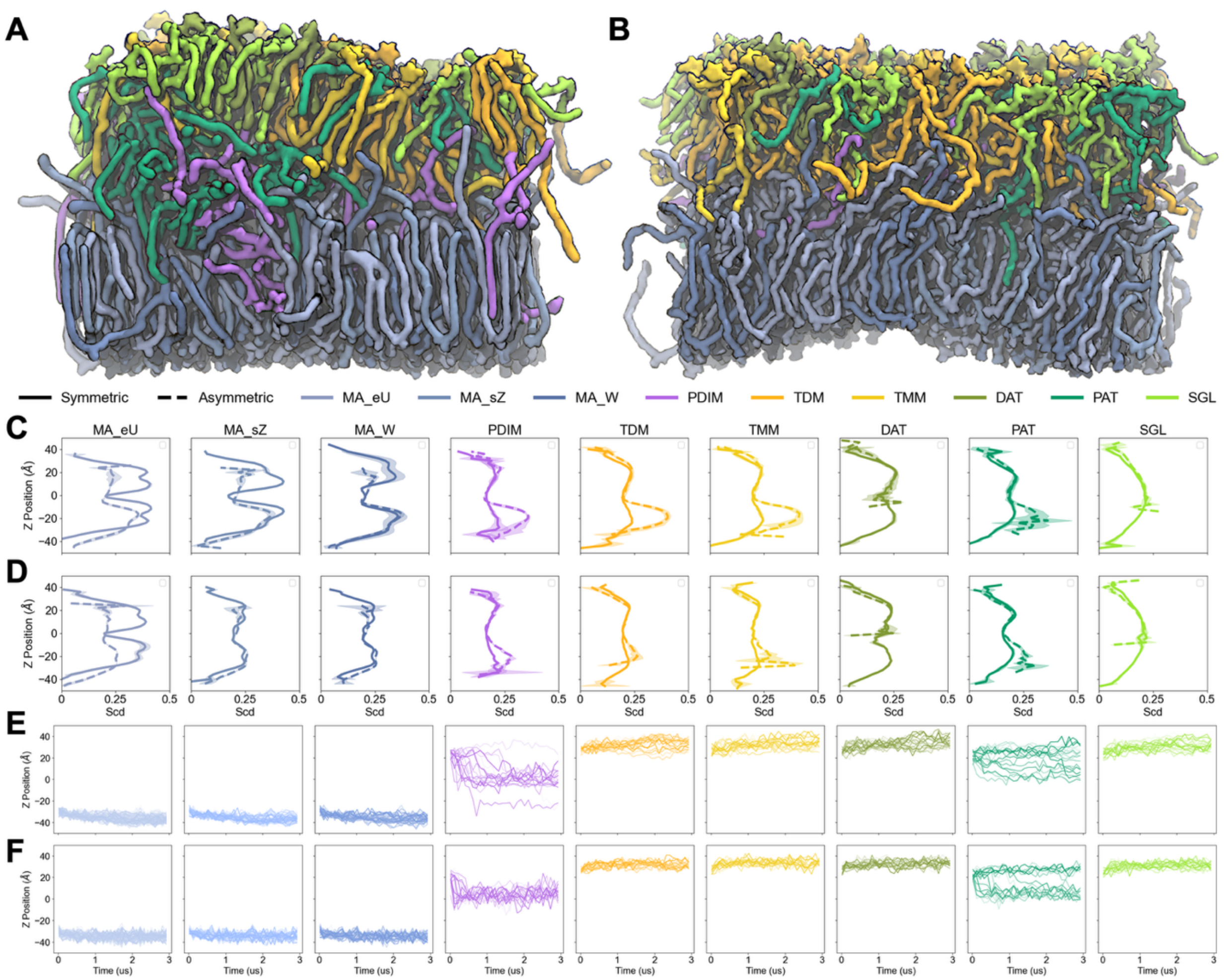


**Figure S7. Scd & Headgroup Time Series for Asymmetric Systems. A,B.** Snapshots after 3 μs of production of asymmetric bilayers at 313 K and 333 K, respectively. **C,D.** Comparison of the average deuterium order parameter for each lipid type along the z-axis for symmetric (solid lines) and asymmetric (dashed lines) bilayers at 313 K and 333 K, respectively. **E,F.** Headgroup z position time series for each lipid type in the asymmetric systems at 31 3K and 333 K, respectively.

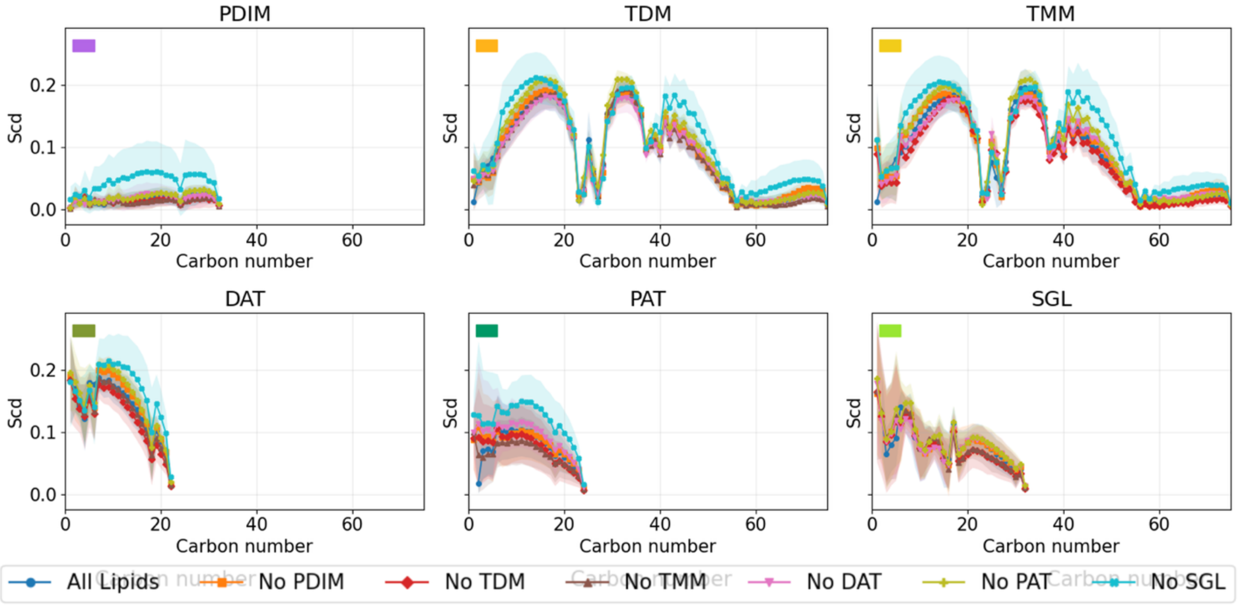


**Figure S8. Scd vs Carbon Number for Outer Leaflet Symmetric Bilayers.** All results shown are for systems simulated at 313K.
